# Predicting identity-preserving object transformations across the human ventral visual stream

**DOI:** 10.1101/2020.08.13.250191

**Authors:** Viola Mocz, Maryam Vaziri-Pashkam, Marvin Chun, Yaoda Xu

## Abstract

In everyday life, we have no trouble recognizing and categorizing objects as they change in position, size, and orientation in our visual fields. This phenomenon is known as object invariance. Previous fMRI research suggests that higher-level object processing regions in the human lateral occipital cortex may link object responses from different affine states (i.e. size and viewpoint) through a general linear mapping function with the learned mapping capable of predicting responses of novel objects. In this study, we extended this approach to examine the mapping for both Euclidean (e.g. position and size) and non-Euclidean (e.g. image statistics and spatial frequency) transformations across the human ventral visual processing hierarchy, including areas V1, V2, V3, V4, ventral occipitotemporal cortex (VOT), and lateral occipitotemporal cortex (LOT). The predicted pattern generated from a linear mapping could capture a significant amount, but not all, of the variance of the true pattern across the ventral visual pathway. The derived linear mapping functions were not entirely category independent as performance was better for the categories included in the training. Moreover, prediction performance was not consistently better in higher than lower visual regions, nor were there notable differences between Euclidean and non-Euclidean transformations. Together, these findings demonstrate a near-orthogonal representation of object identity and non-identity features throughout the human ventral visual processing pathway, with the non-identity features largely untangled from the identity features early in the visual processing.

**Significance Statement:** Presently we still do not fully understand how object identity and non-identity (e.g. position, size) information are simultaneously represented in the primate ventral visual system to form invariant representations. Previous work suggests that the human lateral occipital cortex may be linking different affine states of object representations through general linear mapping functions. Here we show that across the entire human ventral processing pathway, we could link object responses in different states of non-identity transformations through linear mapping functions for both Euclidean and non-Euclidean transformations. These mapping functions are not identity-independent, suggesting that object identity and non-identity features are represented in a near, rather than a completely, orthogonal manner.

## Introduction

Even though objects are constantly changing in position, size, and orientation in our visual fields as we move through the world, we can keep track of their identities. Such invariant object identity representations are thought to be achieved through visual processing in the primate ventral visual pathway, where tangled object identity representations become linearly separable and transformation tolerant from lower to higher visual regions (e.g., DiCarlo & Cox, 2007; Isik et al., 2013; Rolls, 2000; Rust & DiCarlo, 2010; see the illustration in Figure 1A).

**Figure 1.**
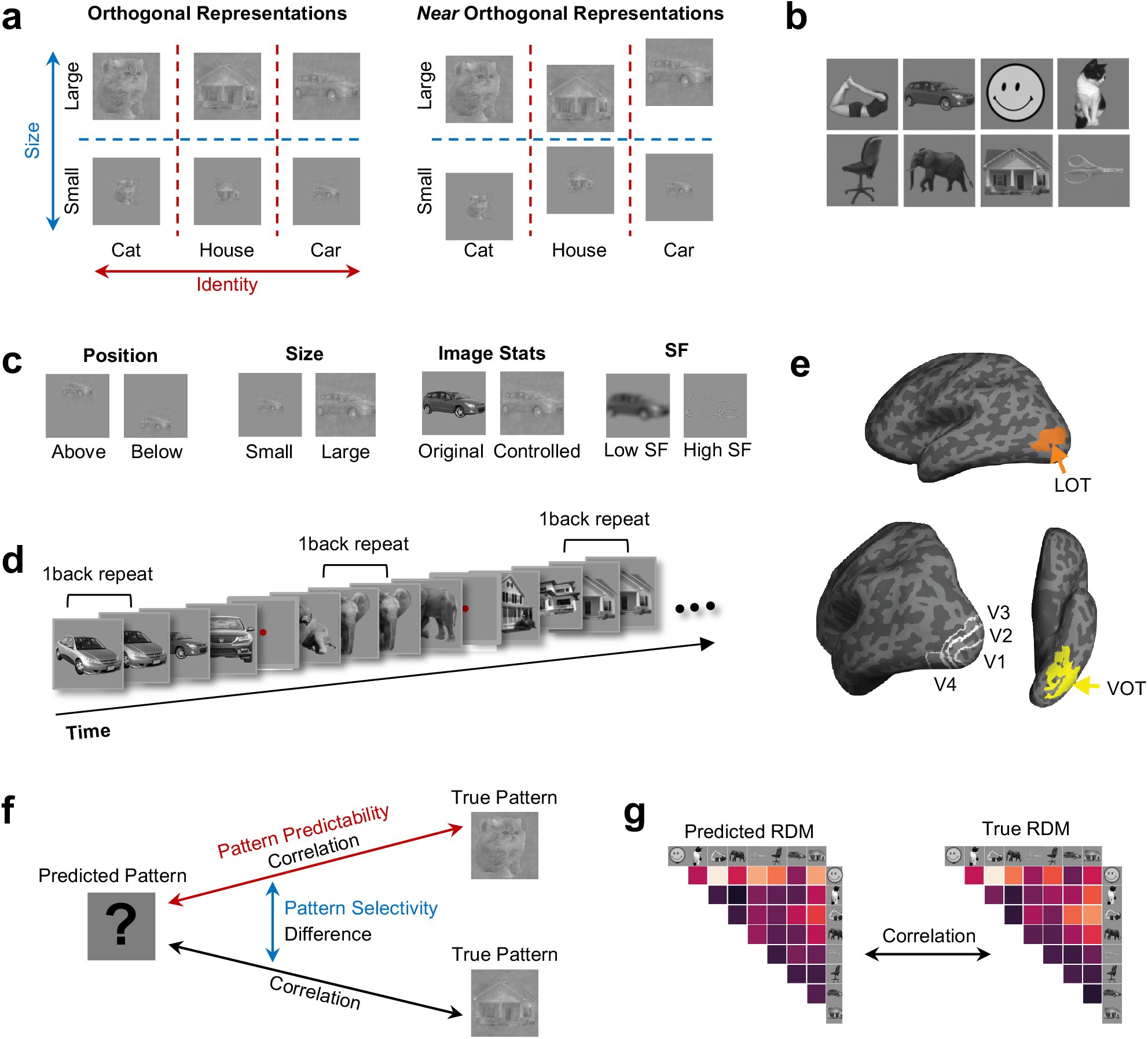
Possible neural representational space structures, experimental details, and analyses used. **a.** A schematic illustration of how object identity and non-identity features may be represented together in high-dimensional neural representational space, using size as a non-identity feature. *Left panel*, completely orthogonal representations of these two types of features, with the object responses across the two states of the size transformation being equidistant for each object in representational space. *Right panel*, near orthogonal representations of these two types of features, with the object responses across the two states of the size transformation for each object being different in the representational space. **b.** The 8 natural object categories used. Each category contained 10 different exemplars varying in identity, pose (for the animate categories only), and viewing angle to minimize the low-level image similarities among them. **c.** The four types of non-identity transformations examined, including position, size, image stats, and spatial frequency. Each transformation included two states. **d.** An illustration of the block design paradigm used. Participants performed a one-back repetition detection task on the images. An actual block in the experiment contained 10 images with two repetitions per block. **e.** Inflated brain surfaces from a representative participant showing the ROIs examined. They included topographically defined early visual areas V1 to V4 and functionally defined higher object processing regions LOT and VOT. **f-g.** Illustrations of the analyses performed to evaluate the predicted pattern. **f.** To evaluate pattern predictability, the predicted and the true patterns were directly correlated; to evaluate pattern selectivity, the correlation between the predicted and the true patterns from the same category was compared to the correlation between the predicted and the true patterns from different categories. **g.** To evaluate the preservation of the category representational structure, the representational dissimilarity matrix (RDM) derived from the predicted patterns was vectorized and correlated with that from the true patterns. See Methods for more details.

At the same time, we can keep track of non-identity features such as size, orientation, and position, which are important for tasks like object grasping. Different states of a non-identity transformation, such as the two positions in a position transformation, can also be decoded across object identities in both lower and higher ventral visual processing regions (Hung et al., 2005; Hong et al., 2016; Vaziri-Pashkam et al., 2019). This suggests that object identity and non-identity information may be represented in an orthogonal manner, which would facilitate independent access to these two types of information in different tasks. If so, we should be able to learn a linear mapping between the neural responses to two states of one object, and apply that mapping to a new object’s neural response in one state to predict its neural response in the other state. Indeed, in human lateral occipital cortex (LOC), a higher ventral visual processing region, a linear mapping function has been shown to link different affine states of object representations, even for objects not included in training (Ward et al., 2018).

Presently, it is unknown whether linear mapping predicts object responses across transformations only in higher-level visual processing regions or also in lower-level regions. An object feature untangling view may predict the existence of a stronger linear mapping in higher than lower visual regions as different object features become more separated throughout the ventral visual processing pathway. However, the separation of some non-identity features could occur early during visual processing. Thus, a linear mapping function would similarly predict responses in higher and lower visual areas.

An orthogonal representation would predict that a linear mapping function derived from one object would equally well predict the neural response patterns of that object and other objects after transformation (Figure 1A left). However, the generalization in prediction could be incomplete, suggesting a near-orthogonal representation, such that both an object independent and an object dependent component would be needed to fully capture object responses between two states of a non-identity transformation (Figure 1A right).

In the present study, we attempted to provide answers to these questions. We used existing data from two studies (Vaziri-Pashkam & Xu, 2019; Vaziri-Pashkam et al., 2019) in our analysis. Specifically, we adapted the methodology of Ward et al. (2018), examining the linear mapping between fMRI representations of objects in different states of non-identity transformations in human occipito-temporal cortex (OTC) as well as in topographically defined early visual areas V1-V4. We aimed to determine how identity and non-identity features are simultaneously represented throughout the human ventral visual hierarchy. Taking into account the response reliability of different visual regions, we examined neural responses to two Euclidean transformations (i.e., position and size) and two non-Euclidean transformations (i.e., changes in image statistics and spatial frequency). Besides testing the success of the predicted fMRI response patterns, we also tested how well object representational structure could be preserved through linear mapping using representational similarity analysis (RSA) (Kriegeskorte & Kievit, 2013). We further tested if similarity between objects plays a role in predicting response patterns across objects after linear mapping.

We found that a linear mapping could successfully link object responses in different states of non-identity transformations throughout the human ventral visual stream for both Euclidean and non-Euclidean transformations. However, these mapping functions were not entirely identity-independent, suggesting that object identity and non-identity features are represented in a near-, rather than complete-, orthogonal manner.

## Materials and Methods

In this study, we applied the representational transformation analysis developed by Ward et al. (2018). This analysis method was originally used to examine fMRI responses from objects undergoing affine transformation in the human LOC. For the current study, we used this analysis to examine such responses throughout the entire human ventral processing pathway. In addition to affine/Euclidean transformations studied before, here we documented responses from two Euclidean (position and size) and two non-Euclidean transformations (image stats and SF). To increase SNR, instead of using an event-related design as in Ward et al., we used a block design and examined responses from the average of multiple exemplars of an object category (i.e., category response) rather than individual objects. In Ward et al., the two states of an object were presented one after another within the same trial, raising the possibility that noise correlation between the two states of the object could contribute to the prediction success.

That is, noise from one state of the object could be carried forward to the predicted pattern after linear mapping, making the predicted pattern more similar to the true pattern of the other state of the same object due to temporal proximity than it would otherwise be. By using the block design, we removed such temporal response correlations in the present study. The overall correlations between the predicted and the true patterns were fairly low in Ward et al. (with the mean correlations being less than .15 for Fisher-Z transformed correlation coefficients), raising the possibility that a linear mapping function may only capture a very small amount of variance associated with response changes between two states of an object. Such a low correlation, however, could be due to the event-related design used. By taking into account the response reliability of a brain region in this study, we would be able to evaluate how good the predicted patterns are compared to the true pattern.

During each run of the experiments, human participants viewed blocks of images, each containing 10 exemplars from one of 6 or 8 real-world object categories (faces, houses, bodies, cats, elephants, cars, chairs, and scissors), and performed a 1-back repetition detection task. During our data analysis, we first selected the 75 most reliable voxels from each ROI to equate the number of voxels across ROIs and to increase power (Tarhan & Konkle, 2019; similar results were obtained when all voxels from each ROI were included, see Figure 2-1 for pattern predictability results). We then extracted the fMRI response pattern corresponding to each image block in each run and used that as the fMRI response pattern for the specific object category shown in that block of that run. We used a split-half approach in our analysis and divided the runs into odd and even halves. Within each half, using a leave-one-run-out cross-validation procedure, we derived a linear transformation matrix following Ward et al. (2018) to link the fMRI response patterns of two states of an object category in a given transformation in the training data. Using this transformation matrix, in the test data, we generated the predicted pattern of an object category in one state using its true pattern from the other state.

We evaluated the predicted patterns in three different ways: how well they correlated with the true patterns, whether they showed category selectivity and were more similar to the true patterns of the same than different categories, and whether the category representational structure was preserved among the predicted patterns. In all three analyses, we compared the performance of the predicted patterns to that of the true patterns to evaluate whether a linear mapping can fully capture all the variance associated with the true patterns. We also examined the effect of training category (i.e., whether or not a category was included in the training data), the effect of training set size (i.e., the number of categories included in the training data), the effect of ROI (i.e., whether or not the effect differed among the different brain regions), and their interactions. We additionally examined how the ability of using the response of one category to predict that of another category was determined by the similarity of these two categories in a given brain region.

The details of the four fMRI experiments included are described in two previous publications (Vaziri-Pashkam et al., 2019; Vaziri-Pashkam & Xu, 2019). They are summarized here for the readers’ convenience.

### Participants

Seven (four females), seven (four females), six (four females), and ten (five females) healthy human participants with normal or corrected to normal visual acuity, all right-handed, and aged between 18 and 35 took part in Experiments 1 to 4, respectively. All participants gave their informed consent before the experiments and received payment for their participation. The experiments were approved by the Committee on the Use of Human Subjects at Harvard University.

### Experimental Design and Procedures

In all four experiments, participants performed a 1-back object repetition detection task while viewing blocks of grayscale images of real-world object categories (Figure 1B-1D). Experiments 1 to 3 included 8 real-world object categories and they were: body, car, cat, chair, elephant, face, house, and scissors. Experiment 4 included 6 real-world object categories and they were: body, car, chair, elephant, face, and house. These sets of categories cover a broad range of real-world objects and include small/large, animate/inanimate, and natural/man-made objects. Similar sets have been used in previous investigations of object category representations in the OTC (e.g., Haxby, 2001; Kriegeskorte et al., 2008). Each category contained 10 exemplars that varied in identity, pose (for the animal and body categories only) and viewing angle to minimize the low-level similarities among them. All images were placed on a dark gray square and displayed on a light gray background.

We analyzed two Euclidean and two non-Euclidean transformations. The two Euclidean transformations were: (1) Experiment 1 - position: object image appearing above vs. below fixation, (2) Experiment 2 - size: object image shown in small vs. large size. The two non-Euclidean transformations were: (1) Experiment 3 - image stats: object images shown in original vs. controlled format, and (2) Experiment 4 - Spatial Frequency (SF): object images shown in high vs. low SF content of an image (Figure 1C). Controlled images were generated using the SHINE technique to achieve spectrum, histogram, and intensity normalization and equalization (Willenbockel et al., 2010). Controlled images also appeared in Experiments 1 and 2 to better equate low-level differences among the images from the different categories.

During the experiment, blocks of images were shown. Each block contained a random sequential presentation of 10 exemplars from the same object category and the same transformation condition (e.g. for Experiment 1, in one block of a run, all the images were of cats positioned above fixation). Each image was presented for 200 ms followed by a 600 ms blank interval between the images (Figure 1D). Two image repetitions occurred randomly in each block. Participants were asked to view the images and report the repetitions by pressing a key on an MR-compatible button box. To ensure proper fixation, participants fixated at a central dot throughout the experiment, and eye movements were monitored in all four experiments using an SR-research Eyelink 1000 eyetracker. Each block lasted 8 sec followed by an 8 sec fixation period. In Experiments 1 to 3, there was an additional fixation period of 8 sec at the beginning of each run. In Experiment 4, there was an additional fixation period of 12 sec at the beginning of each run.

Each run within Experiments 1 to 3 contained 16 blocks, 1 for each of the 8 object categories in each of the 2 states of the transformation. Each run within Experiment 4 contained 18 blocks, 1 for each of the 6 object categories in the low SF condition, the high SF condition, and the full SF condition. The full SF condition blocks were not used in the transformation analysis. Experiments 1 to 3 included 16 runs, with each run lasting 4 min 24 sec. Experiment 4 included 18 runs, with each run lasting 5 min. The order of the object categories and the 2 stages of the transformation were counterbalanced across runs and participants.

### Data Analysis

#### ROI Definitions

We examined ROIs from topographically localized areas within occipital cortex, including V1, V2, V3, and V4 in each participant (Figure 1E, left panel). We also examined the functionally localized regions VOT and LOT (Figure 1E, middle and right panels) (see Vaziri-Pashkam & Xu, 2019 and Vaziri et al., 2019 for details of these localizers and MRI methods). LOT and VOT loosely correspond to the location of lateral occipital (LO) and posterior fusiform (pFs) areas (Grill-Spector et al., 1998; Kourtzi & Kanwisher, 2000; Malach et al., 1995) but extend further into the temporal cortex in an effort to include as many object selective voxel as possible in the OTC. To generate fMRI response patterns for each condition in each run, we first convolved the 8 sec stimulus presentation boxcars with a hemodynamic response function. Then, for Experiments 1 to 3, we conducted a general linear model analysis with 16 factors (2 states of the transformation × 8 object categories) to extract beta value for each condition in each voxel in each ROI. This was done separately for each run. For Experiment 4, we performed a general linear model analysis with 18 factors (3 spatial frequency conditions × 6 object categories) to extract the beta value in a similar way, again separately for each run. While the full SF condition was used in subsequent analysis in Vaziri-Pashkam et al. (2019), it is not used in subsequent analysis here. We z-normalized the beta values across all voxels for each condition in a given ROI in each run to remove amplitude differences between conditions, ROIs and runs.

#### Reliability-based Voxel Selection

As pattern decoding to a large extent depends on the total number of voxels in an ROI, to equate the number of voxels in different ROIs to facilitate comparisons across ROIs and to increase power, we selected 75 most reliable voxels in each ROI using reliability-based voxel selection (Tarhan & Konkle, 2019). Across the experiments, the ROIs ranged from 115 to 901 voxels before voxel selection. This method selects voxels whose response profiles are consistent across odd and even halves of the runs and works well when there are around 15 conditions. To implement this method, for each voxel, we calculated the split-half reliability by first averaging the runs within the odd and even halves and then correlating the resulting averaged responses for all conditions (12 or 16 in total for the 6 or 8 image categories and 2 states of a transformation) across the even and odd halves. We then selected the top 75 voxels with the highest correlations. To avoid circularity in analysis, these 75 voxels were selected using only the training runs in each iteration of a leave-one-out cross validation procedure (which will be explained in more detail in the next section). The 75 voxels chosen maintained a high split-half reliability of at least *r = 0.70* for each participant and ROI while providing an optimal number of features for subsequent ridge regression analysis. It is important to note, however, that the results remained the same in all experiments when we included all voxels from each ROI in the analysis. Thus the pattern of results we obtained was fairly stable and did not depend on the inclusion of the most reliable voxels. Nevertheless, we report in the main paper results from the most reliable voxels to equate the number of voxels selected from each ROI and to ensure that the results obtained were not due to noise and a lack of power.

#### Deriving Linear Mapping Between Two States of a Transformation

We adapted the linear mapping analysis used by Ward et al. (2018) with a split-half analysis on the 75 most reliable voxels for each training iteration in each ROI. Training and testing of the linear mapping were computed separately for each participant, each ROI, and each state of the transformation in each experiment. For each ROI, the data were first split into even and odd runs. Within each half of the data, a leave-one-out cross-validation procedure was conducted where one run served as the testing run while the remaining runs served as training runs. For Experiments 1 to 3, which had 16 runs, this meant that first the data were split into 2 groups with 8 runs each, and within each group there were 7 training runs and 1 testing run. For Experiment 4, which had 18 runs, this meant that first the data were split into 2 groups with 9 runs each, and within each group there were 8 training runs and 1 testing run. The training runs of the even and odd runs were used to select the top 75 most reliable voxels of an ROI as described in the previous section. These 75 most reliable voxels were then applied to the testing runs. Additionally, for Experiments 1 to 3, the number of object categories included in training ranged from 1 to 7 as these experiments included 8 categories in total. For Experiment 4, the number of object categories included in training ranged from 1 to 5 as this experiment included 6 categories in total. All possible combinations of training categories were used.

During training of each leave-one-out cross-validation fold, we derived linear mapping matrices to link the 2 states of a transformation in both directions (e.g. from small to large and from large to small) using ridge regression. The end result was that responses from the 75 voxels in one state, after convolving with the trained linear mapping matrix, would predict the responses of the 75 voxels in the other state. This was accomplished in the following way. For each object category, we first constructed two 75 [voxels] × (7 or 8) [training folds] matrices corresponding to the original and the transformed states, respectively. If more than one object category was included in training, we concatenated each object’s matrix so that the full pattern matrix was 75 [voxels] × (7 or 8) [training folds] × (1 to 7) [training categories]. Using ridge regression, we derived the linear mapping ***β*** between the original state (matrix ***X***) and the transformed state (matrix ***Y***) as follows:

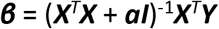

where ***β*** is a 75 [voxels] × 75 [voxels] linear mapping matrix determining how each voxel in one transformation state should be weighted to predict each voxel in the other transformation state, ***ɑ*** is the ridge regression penalty (set as ***ɑ*** = 1), and ***I*** is the identity matrix. Regularizing the linear model with a penalty is necessary as there were fewer observations [training folds × training categories] than features [voxels]. ***β*** was then applied to the left-out data to predict the response pattern in one transformation state given the response pattern of the other transformation state.

#### Experimental Design and Statistical Analyses

Seven, seven, six, and ten human participants took part in Experiments 1 to 4, respectively. These numbers were chosen based on prior published studies (e.g., Haxby et al., 2001; Kamitani & Tong, 2005). The factors described in the remaining methods sections were evaluated at the group level using repeated-measures ANOVA and post-hoc t-tests. We corrected for multiple comparisons in all post-hoc analyses using the Benjamini-Hochberg method (Benjamini & Hochberg, 1995). All statistical tests were conducted using R (R Core Team, 2018).

#### Testing the Predictions of the Learned Linear Mapping

To test how well the learned linear mapping could predict fMRI response patterns between the two states of a given transformation, we first generated predicted patterns using the learned linear mapping. We then compared the predicted and true patterns using Pearson correlation. Specifically, within each half of the data, for each leave-one-out cross-validation fold, we first generated two predicted patterns (one for each state) for each object category from the left-out data by using the pattern from one state to predict the other state using the learned linear mapping matrices (one for each direction of transformation). We then correlated each predicted pattern from one half of the data with the corresponding true pattern in the other half of the data (Figure 1F). We normalized the resulting correlation coefficients by dividing them by each ROI’s split-half reliability (method similar to Kietzman et al., 2019).

Because different ROIs likely had different reliability, normalizing the correlation by each ROI’s reliability measure enabled us to directly compare the results across the different ROIs. To derive the reliability of a given ROI, we averaged the runs within the odd and even halves and correlated the corresponding averaged true patterns for each condition across the two halves of the data; we then averaged all the resulting correlations as our measure of split-half reliability for a given ROI. The normalized correlation coefficients were averaged across categories and the different training and testing folds for each participant in each ROI.

The correlation between the predicted and true patterns is a measure of how well a linear mapping captures the change in response patterns between two states of an object. A correlation no different from 1 would show a full capture of the data, a correlation greater than 0 but less than 1 would show a partial capture of the data, and a correlation no different from 0 would show no capture of the data. Besides examining the correlation between the predicted pattern with the corresponding true pattern, we also examined whether such a correlation was higher for the same than the different category pairs. This allowed us to further evaluate category selectivity of the predicted pattern.

Within each transformation type and each ROI, we also assessed the effect of category (i.e., prediction performance for categories included or not included in training) and the effect of training set size (i.e., how prediction performance depended on the number of categories included in the training data) and their interaction. These analyses allowed us to test the generalizability of the learned linear mapping: how well a linear mapping learned from one set of categories could successfully predict the patterns of categories not included in training.

To test if linear mapping better predicts the neural response pattern in higher than lower visual regions, we compared performance amongst the ROIs. To account for the effect of training set size and to streamline the analysis, we only included the lowest and the highest training set sizes from each transformation and tested the effects of ROI, training category, training set size, and their interactions. For each of the two training set sizes and two training categories, we further examined if there was a positive linear trend in performance from lower-level to higher-level visual regions. This was done by fitting a regression line for each participant and then testing whether the resulting slopes were greater than 0 at the group level using a one-tailed t-test.

### Evaluating the Predicted Representational Structure

Using representational similarity analysis (RSA) (Figure 1G) (Kriegeskorte & Kievit, 2013), we tested if the representational structure of category information is preserved for the predicted patterns. To do so, for each transformation state, each half of the data, and each category, we first averaged the predicted patterns across runs. We also averaged the true patterns across runs for each half of the data. We then constructed representational dissimilarity matrices (RDMs) for the predicted patterns as well as the true patterns, with each cell of the matrix corresponding to the one minus the Pearson correlation coefficient of a pair of category patterns. The RDM captures the relative similarity among the different categories. To test if the RDM of the predicted patterns is similar to that of the true patterns, we first vectorized the upper triangle of each RDM to generate a category similarity vector. We then calculated the Pearson correlation of the category similarity vector from the predicted pattern in one half of the data to the true category similarity vector in the other half of the data. The resulting correlation was then averaged across the two halves of the data. This analysis was performed separately for the predicted patterns of trained categories and the predicted patterns of untrained categories across each training set size. To account for differences in measurement reliability across the different ROIs, these correlations were further divided by the split-half reliability of each ROI. To calculate each ROI’s split-half reliability, within each state of a transformation, we calculated the correlation of the true pattern RDM in one half of the runs with the true pattern RDM in the other half. The resulting correlations were averaged between the two states of the transformation to produce the split-half RDM reliability measure for that ROI.

#### Relating Category Similarity to Pattern Prediction Generalization

In this analysis, we examined how the generalizability of linear mapping may depend on the similarity between the different categories in a given brain region. To do so, for each ROI, using the cross-validated split-half analysis described earlier, we first obtained the linear mapping between the two states of a given transformation using data from only one category as the training data to predict the response of all other categories from one state to the other state. We then correlated the predicted pattern and the true pattern of the same category across the two halves of the data as described earlier. The resulting correlation coefficient was used as the prediction score for how well the training data of a given category could successfully predict the pattern of a different category. We repeated this analysis by including the data from each category as the training data to predict the response patterns of all other categories from one state to the other state. The results were averaged between the two states of a transformation and the two directions of predictions (i.e., using X to predict Y and using Y to predict X) and were used to construct a prediction similarity matrix in which each cell of the matrix reflects how well two categories may predict each other. To obtain the similarity between the object categories, we constructed a category similarity matrix with each cell of the matrix being the pairwise correlation of the true patterns of two categories across the two halves of the data. We vectorized the off-diagonal elements of these matrices and correlated them. If the generalizability of transformations depends on the similarity between the different categories in a given brain region, then we expected to see a high correlation between prediction similarity and category similarity. We did not correct the prediction similarity matrix and the category similarity matrix by the split-half reliability of each ROI here, as such correction would not affect the final correlation results (since values were z-normalized during correlation calculation and thus any scaling factor would have no effect).

## Results

Previous research has shown that we can derive linear mappings within human LOC for affine changes of objects that are generalizable to objects not included in training (Ward et al., 2018). We evaluated and extended this work using data from four existing fMRI experiments to test how Euclidean and non-Euclidean object transformations are represented throughout the human ventral visual hierarchy, which included V1 to V4, LOT and VOT. We evaluated the predicted patterns in three different ways: how well they correlated with the true patterns, whether they show category selectivity and were more similar to the true patterns of the same than different categories, and whether the category representational structure was preserved among the predicted patterns. We also examined the effect of training category (i.e., whether or not a category was included in the training data), the effect of training set size (i.e., the number of categories included in the training data), the effect of ROI (i.e., whether or not the effect differed among the different brain regions), and their interactions. We additionally examined how the ability of using the response of one category to predict that of another category was determined by the similarity of these two categories in a given brain region.

### Evaluating the Predicted fMRI Response Patterns

To understand how well a linear function can capture fMRI pattern differences between two states of an object in a given transformation, we evaluated how close the predicted pattern was to the true pattern by correlating the predicted patterns derived from one half of the data with the corresponding true patterns from the other half. To compare across ROIs and to account for differences in data reliability across the different ROIs, we calculated the reliability of the data in each ROI by correlating the corresponding patterns between the two halves of the true data and taking the average of these correlations across categories and the two states of the transformation as the reliability measure. We then normalized the predicted and true pattern correlations by dividing them by the reliability of the data. This was done separately for each ROI in each participant. The results were then evaluated at the participant group level using statistical tests. If the correlation between the predicted and true patterns is no different from 1, it would indicate that the predicted pattern is as good as the true pattern in its correlation with the corresponding pattern in the other half of the experiment. On the other hand, if the correlation is significantly lower than 1, it would indicate that a linear mapping does not fully capture the difference in fMRI response patterns between the two states of a given transformation for an object category.

For all four transformation types and across both the training category and training set size manipulations, we found that normalized predicted and true pattern correlations were overall quite high, ranging between 0.75 and 1, and were all significantly above 0 (see the significance level for each condition indicated by the asterisks on Figure 2; all pairwise comparisons reported here and below were corrected for multiple comparisons using the Benjamini-Hochberg method, see Benjamini & Hochberg, 1995). However, with the exception of three correlations being marginally different from 1, all other correlations were significantly different from 1 across the different ROIs, training categories, training set sizes and transformations (see the significance levels reported on Figure 2A). These results indicate that, for the most part, a linear mapping could capture a significant amount of, but not all, the changes associated with these transformations.

**Figure 2.**
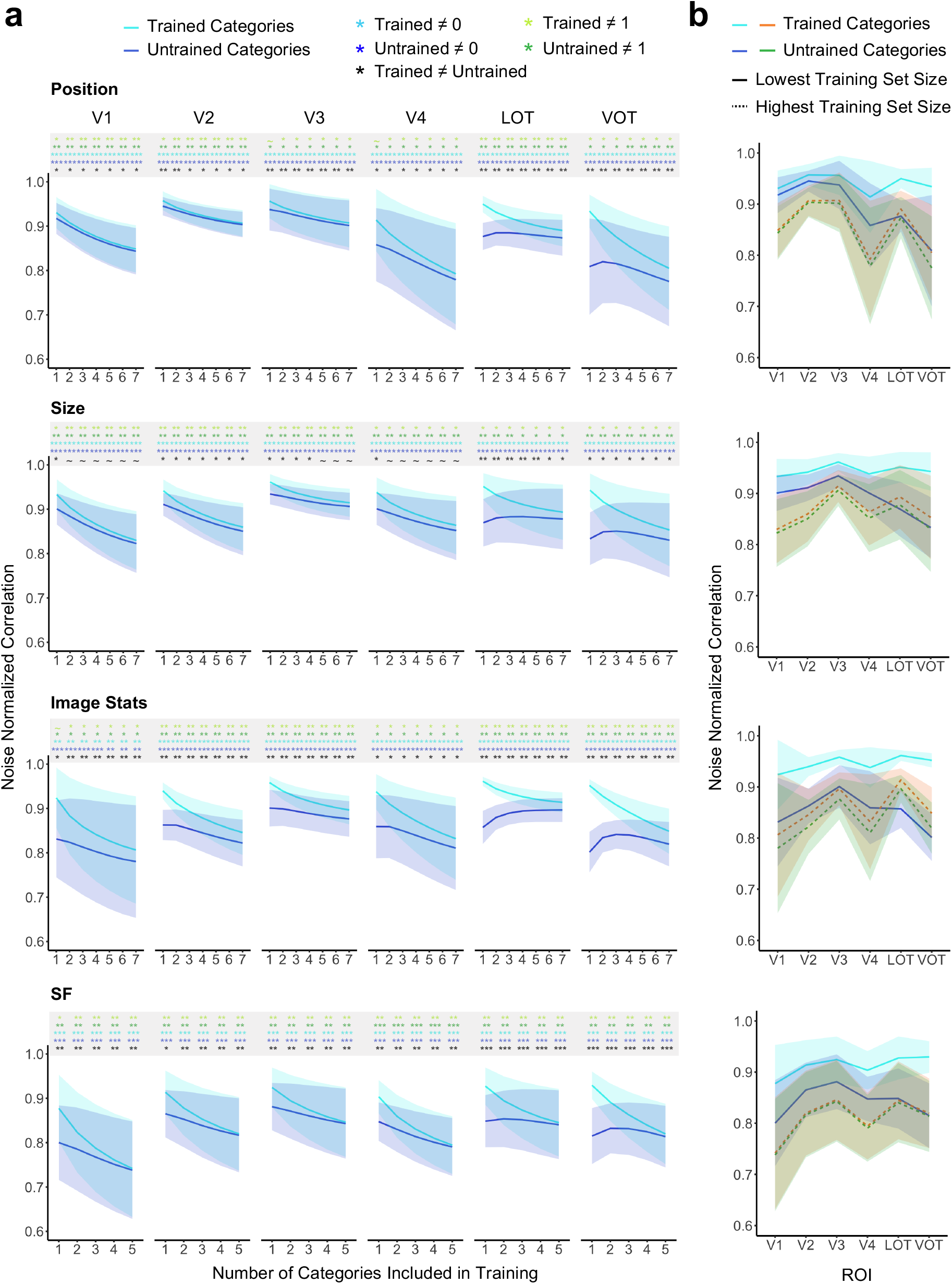
Correlations between the predicted and the corresponding true patterns for each transformation for the 75 most reliable voxels included in each ROI. The correlations were normalized by the reliability of each ROI (see Methods). **a.** Results plotted by ROI and transformation type showing pattern prediction for categories included in the training data (Trained Categories) and those that were not (Untrained Categories) as a function of the number of categories included in the training data. Significant values for pairwise t tests against 0 and 1 for each condition and for the difference between the trained and untrained categories for each training set size are marked with asterisks at the top of each plot. All t tests were corrected for multiple comparison using the Benjamini-Hochberg method. ~ .05 < *p* < .10, * *p* <.05, ** *p* <.01, *** *p* <.001. **b.** Results plotted by transformation type showing pattern prediction as a function of the ROI including only the smallest and the largest training set sizes. Error bars represent the between-subjects 95% confidence interval.

To understand the generalizability of linear mapping in pattern prediction, within each transformation, we next examined in each ROI the effect of training category, training set size, and their interactions. The significance levels of these effects are reported in Table 1. Overall, the main effect of training category was significant in LOT and VOT across all four types of transformations, such that including a category in the training data significantly improved prediction; this effect was less consistent in early visual areas. Post-hoc tests confirmed this and revealed that the effect of training category was present across training set sizes and transformation types, more consistently so in higher than lower visual regions (see the significance levels reported on Figure 2A). Meanwhile, the main effect of training set size produced no benefit across ROIs and transformation types, such that including more categories in the training data either decreased or had no effect on the overall prediction performance (see Figure 3A for a schematic diagram explaining why this may be happening). Interactions between training set size and training category were weak in general, only reaching significance for LOT and VOT for image stats transformation, but absent in all the other conditions.

**Table 1.**
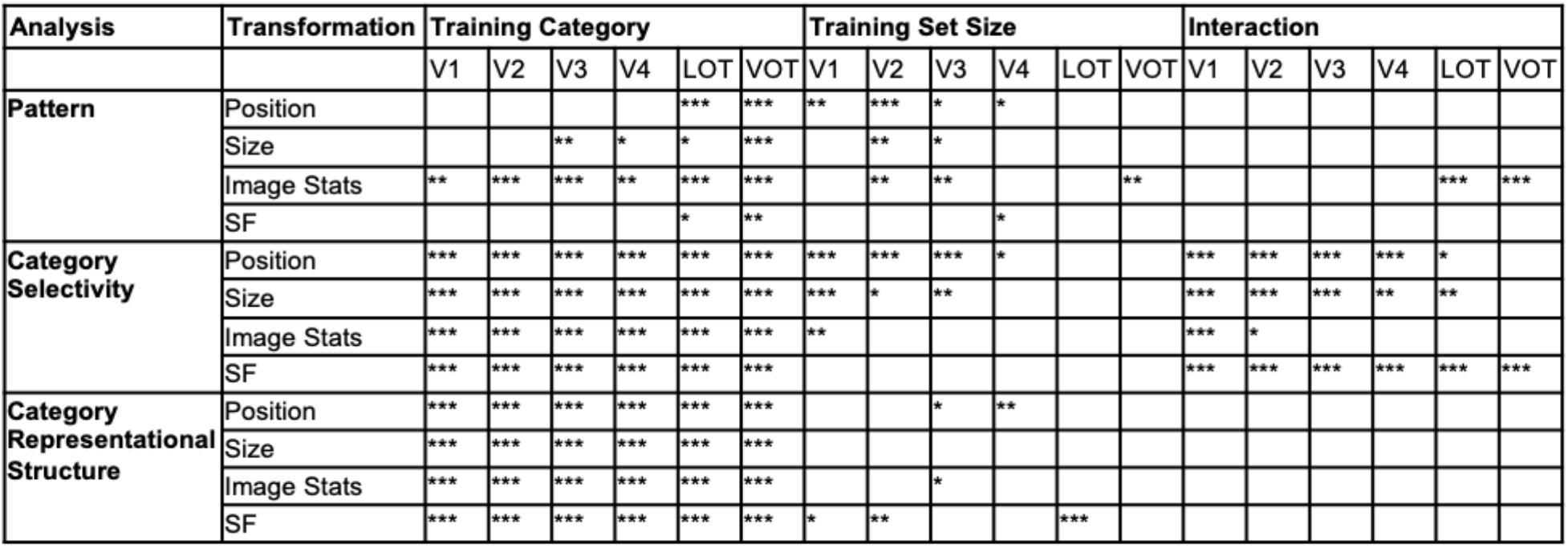
Summary of statistical results of the predicted fMRI response patterns. The predicted patterns were evaluated in terms of their pattern correlation with the true response patterns, their category selectivity (i.e. testing if the predicted pattern was more similar to the true pattern of the same than different categories), and category representational structure. In all three analyses, the effects of training category (i.e., whether or not a category was included in the training data), training set size (i.e., the number of categories included in the training data), and their interactions were examined. The ROIs with significant effects are marked with asterisks. All tests were corrected for multiple comparison using the Benjamini-Hochberg method. * *p* < .05, ** *p* < .01, *** *p* < .001.

**Figure 3.**
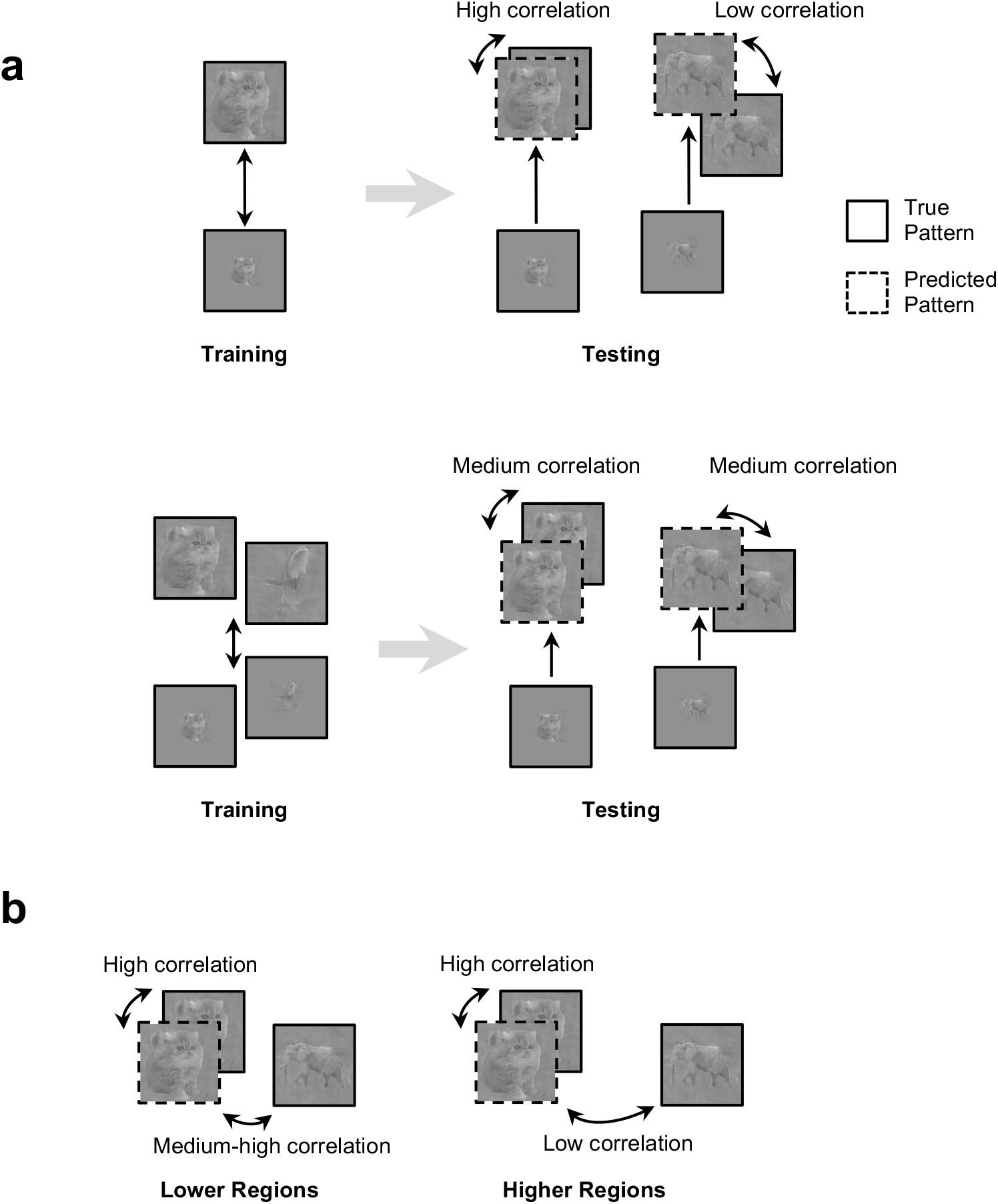
Schematic illustrations of the pattern predictability and pattern selectivity results when object identity and nonidentity features are represented in a near-orthogonal manner. **a.***Top panel*, predictability results of training only one category (e.g. cat). The mapping derived is specific to the one trained category. As such, the correlation between the true and predicted patterns would be higher for the category included in training than that not included in training. *Bottom panel*, predictability results of training two categories. Because the mapping derived is not specific to any one trained category, the correlation between the true and predicted patterns could decrease for the category included in training and increase for the category not included in training. **b.** Predictability and selectivity results in lower and higher visual regions. Because objects are more correlated in lower than higher visual regions to begin with, even when both regions could equally well predict an object category response pattern after training, category selectivity would be greater for higher than lower visual regions. Category selectivity is defined as the difference in the correlation of a predicted pattern with the true pattern of the matching category and the correlation of a predicted pattern with the true pattern of mismatching categories (i.e., is the predicted pattern significantly closer to that of the matching category than mismatching categories?).

To understand if linear mapping better predicts the neural response pattern in higher than lower visual regions, we examined performance amongst the ROIs (Figure 2B). To account for the effect of training set size and to streamline the analysis, we only included the lowest and the highest training set sizes from each transformation and tested the effects of ROI and how it may interact with the effect of training category and training set size. Overall, only image stats transformation showed a main effect of ROI (*F(5,96)* = 2.70, *p* < .05), with pattern prediction being higher for V3 and VOT than the other ROIs (post-hoc paired t tests, *ts* > 2.20, *ps* < .05). No other transformation showed a main effect of ROI (*Fs* < 1.59, *ps* > 0.16). There was also no interaction between the main effect of ROI and that of either training category or training set size (*Fs* < 0.94; *ps* > 0.46). To evaluate whether prediction performance increased from lower to higher visual regions, we tested the existence of a positive linear trend across the ROIs by fitting a regression line across the ROIs in each participant and testing whether the slope was positive at the group level. Across the different transformations and the different training set size and train category combinations, no slope was significantly greater than 0 (*ts* < 1.23, *ps* > 0.21). This suggests that, overall, linear mapping did not better predict the neural responses in higher than lower visual regions across all four types of transformations.

### Evaluating the Selectivity of Predicted fMRI Response Patterns

A successful linear prediction for transformation would not only predict that the predicted patterns are similar to the true patterns, but also that the predicted patterns would be more similar to the true patterns from the same than different categories, thereby showing a high category selectivity. To evaluate this, here we tested whether the difference in correlation between the same and different categories was greater than 0. For the categories included in training, correlation differences for all transformation types, ROIs, and training set sizes were significantly greater than 0 (see the significance levels reported on Figure 4; additionally, see Figure 4-1 for similar results when all voxels from each ROI were included in analysis). For categories not included in training, for all transformation types, correlation differences in LOT and VOT tended to be significantly (or marginally significantly) greater than 0 for large training set sizes; the effect was less consistent for small training set sizes. Correlation differences in V1 to V4 in general did not exceed 0, except in a few cases for the SF and image stats transformations (see the significance levels reported on Figure 4A). These results showed that, when a category was included in the training data, category selectivity was preserved in the predicted patterns across all ROIs, training set sizes, and transformations. However, when a category was not included in the training data, while category selectivity was still preserved in the predicted patterns for higher visual regions, especially when more categories were included in the training, its preservation in lower visual regions was less consistent.

**Figure 4.**
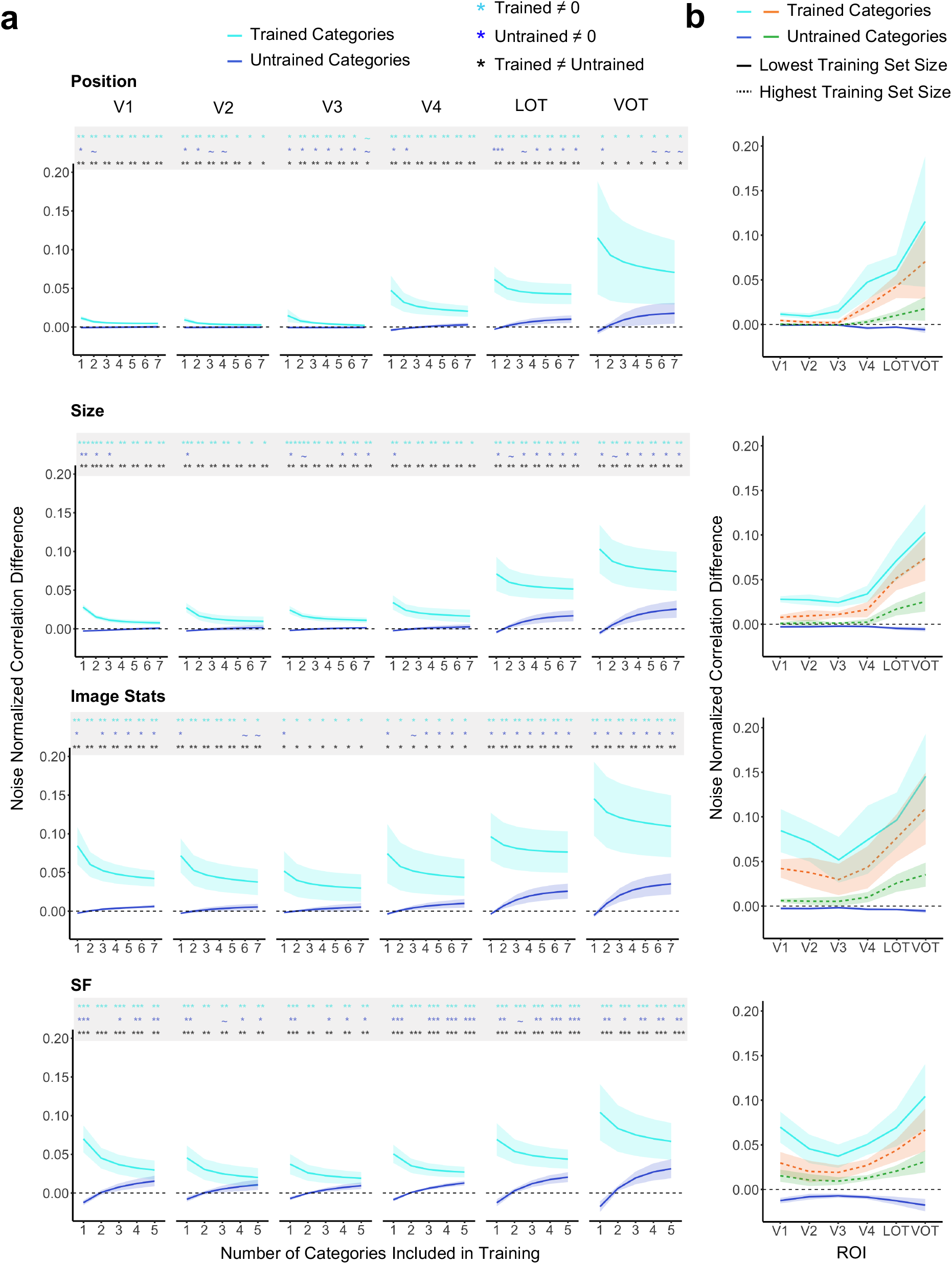
Category selectivity for the predicted patterns for the 75 most reliable voxels included in each ROI. Category selectivity was calculated as the correlation difference between the predicted and true patterns of the same category and that of different categories. **a.** Results plotted by ROI and transformation type showing category selectivity for categories included in the training data (Trained Categories) and those that were not (Untrained Categories) as a function of the number of categories included in the training data. Significant values for pairwise t tests against 0 for each condition and for the difference between the trained and untrained categories for each training set size are marked with asterisks at the top of each plot. All t tests were corrected for multiple comparison using the Benjamini-Hochberg method. ~ .05 < *p* < .10, * *p* <.05, ** *p* <.01, *** *p* <.001. **b.** Results plotted by transformation type showing category selectivity as a function of the ROI including only the smallest and the largest training set sizes. Error bars represent the between-subjects 95% confidence interval.

In further analysis, to better understand the generalizability of linear mapping in preserving category selectivity, within each transformation, we examined in each ROI the effect of training category, training set size, and their interactions. The significance levels of the effects are reported in Table 1. Overall, there was a main effect of training category in all ROIs and all transformation types, such that including a category in training significantly improved category selectivity. Post-hoc tests revealed that the effect of training category was present for most of the training set sizes across ROIs and transformations (see the significance levels reported on Figure 4A). The main effect of training set size was present in only some of the lower-level visual areas, with the effect being inconsistent across the different transformation types. The interaction between the main effects of training category and training set size was present in most ROIs and transformation types, such that the difference between categories included and those not included in the training data tended to decrease with increasing training set size. For the categories included in the training data, the main effect of training set size was significant in V1-V4 for all the transformation types (*F*_*s*_ > 2.70, *ps* < 0.05), reflecting the fact that performance tended to decrease with increasing training set sizes; the opposite, however, was found for the categories not included in the training data, such that there was a main effect of ROI for all regions in the SF and image stats transformation, and for 4 out of 6 regions in the size and position transformations (*F*_*s*_ > 2.99, *ps* < 0.05), where there was an increase in performance with increasing training set sizes. These results are consistent with the linear mapping functions being more tailored to the categories included in training, rather than being independent of these categories.

To understand if category selectivity from the predicted patterns was better preserved in higher than lower visual regions, we examined performance amongst the ROIs (Figure 4B). As before, we only included the lowest and highest possible training set sizes and tested the main effect of ROI and its interaction with the main effects of training category and training set size. Overall, all transformations showed a main effect of ROI (*F*_*s*_ > 2.90, *ps* < 0.05), with LOT and VOT showing higher category selectivity than the other regions in each transformation (post-hoc pairwise tests, *ts* > 2.40, *ps* < 0.05). The main effect of ROI had a significant interaction with training category for the position and size transformations (*F*_*s*_ > 4.60, *ps* < 0.001), with the effect of training category being smaller for lower than higher visual regions. No other interaction with the effect of ROI reached significance. Consistent with these observations, fitting a regression line across the ROIs for each participant revealed a significantly positive linear trend of increasing performance from lower-level to higher-level visual regions for 3 out of the 4 training category and training set size combinations across all the transformations (*ts* > 2.28, *ps* < 0.05; see Figure 4B). The exception to this was for categories not included in training in the lowest training set size, where the slopes of the lines were less than 0 for all the transformation types (*ts* < −1.84, *ps* < 0.05). Note that in this condition, category selectivities were largely no different or significantly less than 0, thus showing a lack of success in pattern prediction likely due to the linear mapping function being much more tailored to the training category when only one category was included in the training data. These results show that, on the whole, for predicted patterns that did exhibit category selectivity, category selectivity was greater for higher than lower visual regions. Nevertheless, these results may not be taken to indicate that the linear mapping function better predicted responses in higher than lower visual regions. This is because object category representation has been shown to be much stronger in higher than lower visual regions by previous fMRI decoding studies (e.g., Vaziri-Pashkam & Xu, 2017 & 2019; Vaziri-Pashkam et al., 2019). As such, a greater category selectivity was expected in higher than lower visual regions even if linear mapping function predicted responses equally well in both regions. For a schematic explanation for how pattern predictability can be roughly equal in lower- and higher-level regions while pattern selectivity is better in higher-level regions, see Figure 3B. Although this analysis does not provide us with a clear differentiation between brain regions, we included it here for completeness as it was the main analysis used in Ward et al. (2018).

### Evaluating the Prediction of Representational Structure

In this analysis, we tested if the category representational structure was preserved in the patterns predicted by a linear mapping function. Although the analyses performed so far demonstrated that the predicted patterns had high correlation with the true patterns and showed high category selectivity, whether or not they retained the original category representational structure remains unknown. To test this, we generated a category RDM that included all pairwise correlations of the predicted patterns for each state of a given transformation for each half of the data. We then correlated the category RDM of the predicted pattern from one half of the data with the true category RDM from the other half to test how well category representational structures were preserved. To compare across ROIs and to account for differences in data reliability across the different ROIs, we obtained an RDM reliability measure for each ROI calculated as the correlation of the true category RDMs from the two halves of the data. We normalized the predicted and true category RDM correlations by dividing them by the reliability of the category RDM correlation. This was done separately for each ROI in each participant. The results were then evaluated at the participant group level using statistical tests. Similar to evaluating the prediction of fMRI response patterns, if the correlation between the RDMs of the predicted and true patterns is no different from 1, it would indicate that the representational structure of the predicted pattern is as good as the representational structure of the true pattern in its correlation with the corresponding RDM in the other half of the data. However, if the correlation is significantly lower than 1, it would indicate that a linear mapping function is not able to fully capture the relative similarities among the categories after objects undergo a transformation.

Across all transformation types and ROIs, the normalized RDM correlations for categories included in training were not significantly different from 1 (*ts* < 2.50, *ps* > .05) and all were significantly above 0 (*ts* > 2.60, *ps* < 0.05; the significance levels are reported on Figure 5A; additionally, see Figure 5-1 for similar results when all voxels from each ROI were included in analysis). However, for categories not included in training, across all four transformations, only LOT and VOT had RDM correlations that were significantly above 0 (*ts* > 2.60, *ps* < 0.05), but less than 1 (*ts* > 2.50, *ps* < 0.05). The correlations for V1-V4 were only consistently above 0 for the image stats transformation (*ts* > 2.40, *ps* < 0.05), but not for the other transformations (*ts* < 1.78, *ps* > 0.11). These results showed that, for categories included in training, regardless of the transformation type, category structure was fully preserved in the predicted patterns in both lower and higher visual regions; however, for categories not included in training, such preservation was only found in higher visual regions and largely absent in lower visual regions.

**Figure 5.**
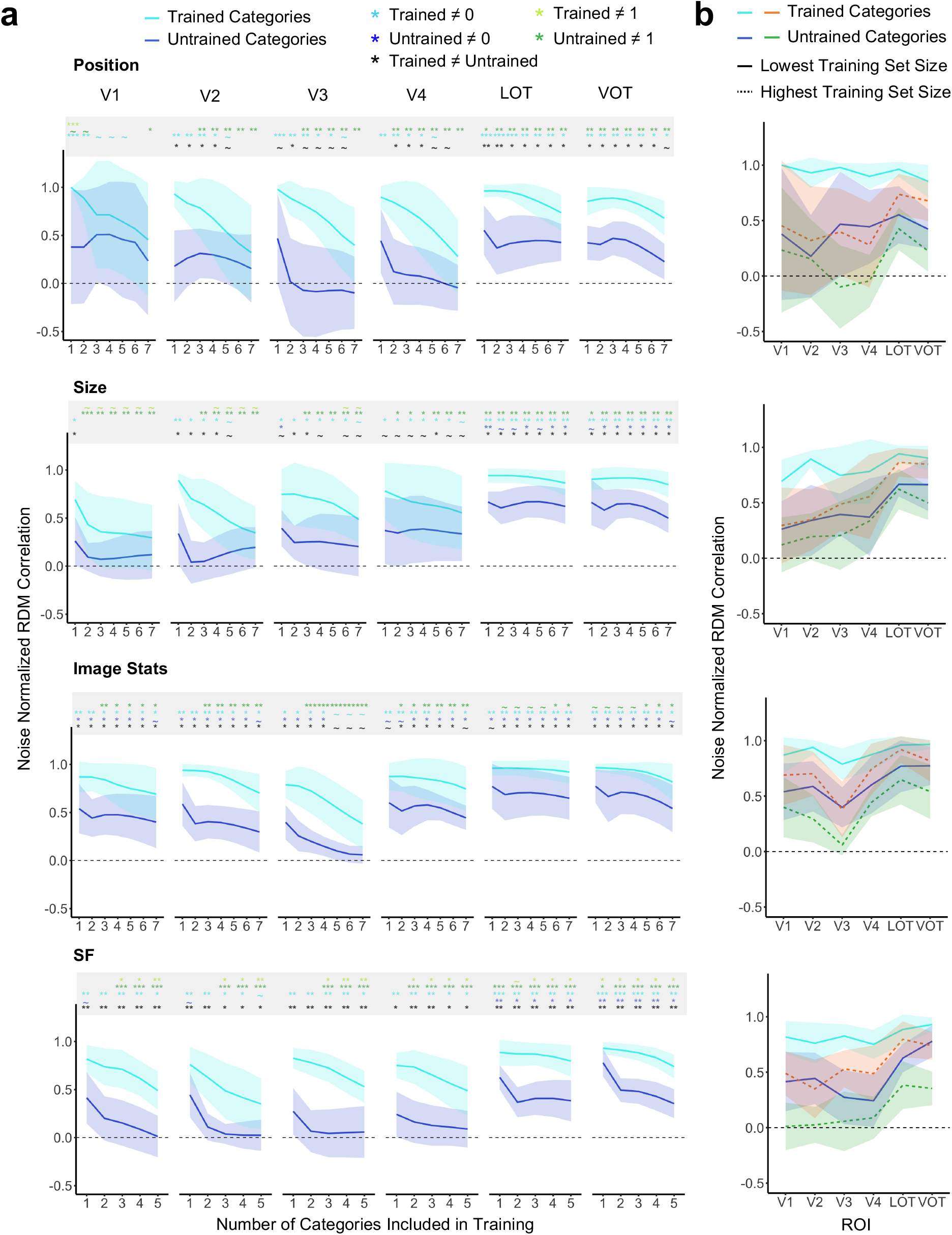
Category RDM correlation between the predicted and true patterns for the 75 most reliable voxels included in each ROI. The correlations were normalized by the reliability of each ROI (see Methods). **a.** Results plotted by ROI and transformation type showing category structure preservation for categories included in the training data (Trained Categories) and those that were not (Untrained Categories) as a function of the number of categories included in the training data. Significant values for pairwise t tests against 0 and 1 for each condition and for the difference between the trained and untrained categories for each training set size are marked with asterisks at the top of each plot. All t tests were corrected for multiple comparison using the Benjamini-Hochberg method. ~ .05 < *p* < .10, * *p* <.05, ** *p* <.01, *** *p* <.001. **b.** Results plotted by transformation type showing category structure preservation as a function of the ROI including only the smallest and the largest training set sizes. Error bars represent the between-subjects 95% confidence interval.

In further analysis, to better understand the generalizability of linear mapping in preserving category representational structure, within each transformation and each ROI, we examined the effect of training category, training set size, and their interactions. The significance levels of the effects are reported in Table 1. There was a main effect of training category in all ROIs and all transformations, with categories included in training showing higher RDM correlations than those not included in training. There were few main effects of training set size and these effects did not follow a consistent pattern across ROIs and transformation types. When there was a main effect, RDM correlations decreased with more categories included in training. There were no interactions between the main effects of training category and training set size.

To understand if category representational structure from the predicted patterns was better preserved in higher than lower visual regions, we examined performance amongst the ROIs (Figure 5B). As before, we only included the lowest and highest possible training set sizes to streamline the analysis and tested the main effects of ROI and its interaction with training category and training set size. A main effect of ROI was found for the size and image stats transformations (*F*_*s*_ > 3.70, *ps* < 0.01), with LOT and VOT showing consistently greater RDM predictions than the other ROIs (post hoc pairwise t-tests, *ts* > 3.50, *ps* < 0.05). The main effect of ROI was absent for the other two types of transformations (*F*_*s*_ < 3.20, *ps* > 0.9). To quantify whether performance increased from lower to higher visual regions, we tested the existence of any positive linear trend across the ROIs in each of the four conditions included in each transformation. Across the 16 total conditions, 10 showed a significant or marginally significant positive trend (*ts* > 1.76, *ps* < 0.093). The exceptions to this were in the position transformation, in which no condition showed a significant trend (*ts* < 1.90, *ps* > 0.87) and in image stats and SF transformations in the lowest training set size for categories included in training (*ts* < 1.29, *ps* > 0.11). Overall, a little over half of the conditions showed better preservation of the representational structure in higher than lower visual regions. There were no interactions between the main effect of ROI and that of either training category or training set size.

### Relating Category Similarity to Pattern Prediction Generalization

Results obtained so far showed that while linear mapping could predict to a great extent fMRI response patterns for object categories in one state of a transformation using responses from another state, the prediction tended to depend, to a significant extent, on the specific categories included in training, rather than being completely independent of the categories included in training. In this analysis, we examined whether categories that are similarly represented in a given ROI could also better predict each other’s response patterns with linear mapping. That is, if category A is more similar to category B than C in a brain region, would A and B show similar linear separation across a transformation such that the linear mapping function derived from the data from category A makes a better prediction for category B than C (see Figure 6A for the two possible scenarios)? To evaluate this possibility, in each ROI, using the same linear mapping procedure, we used data from one category to predict the responses of all other categories from one transformation state to the other state. We then correlated the predicted pattern and the true pattern of the same category across the two halves of the data. This analysis was rotated across all the categories with each serving as the training category to predict the other categories. The results were averaged between the two states of a transformation and the two directions of predictions (i.e., using A to predict B and using B to predict A) and were used to construct a prediction similarity matrix in which each cell of the matrix reflects how well two categories may predict each other. To obtain the similarity between the object categories, we constructed a category similarity matrix with each cell of the matrix being the pairwise correlation of the true patterns of two categories across the two halves of the data within the same state of the transformation averaged over the two states. We vectorized the off-diagonal elements of these matrices and correlated them.

**Figure 6.**
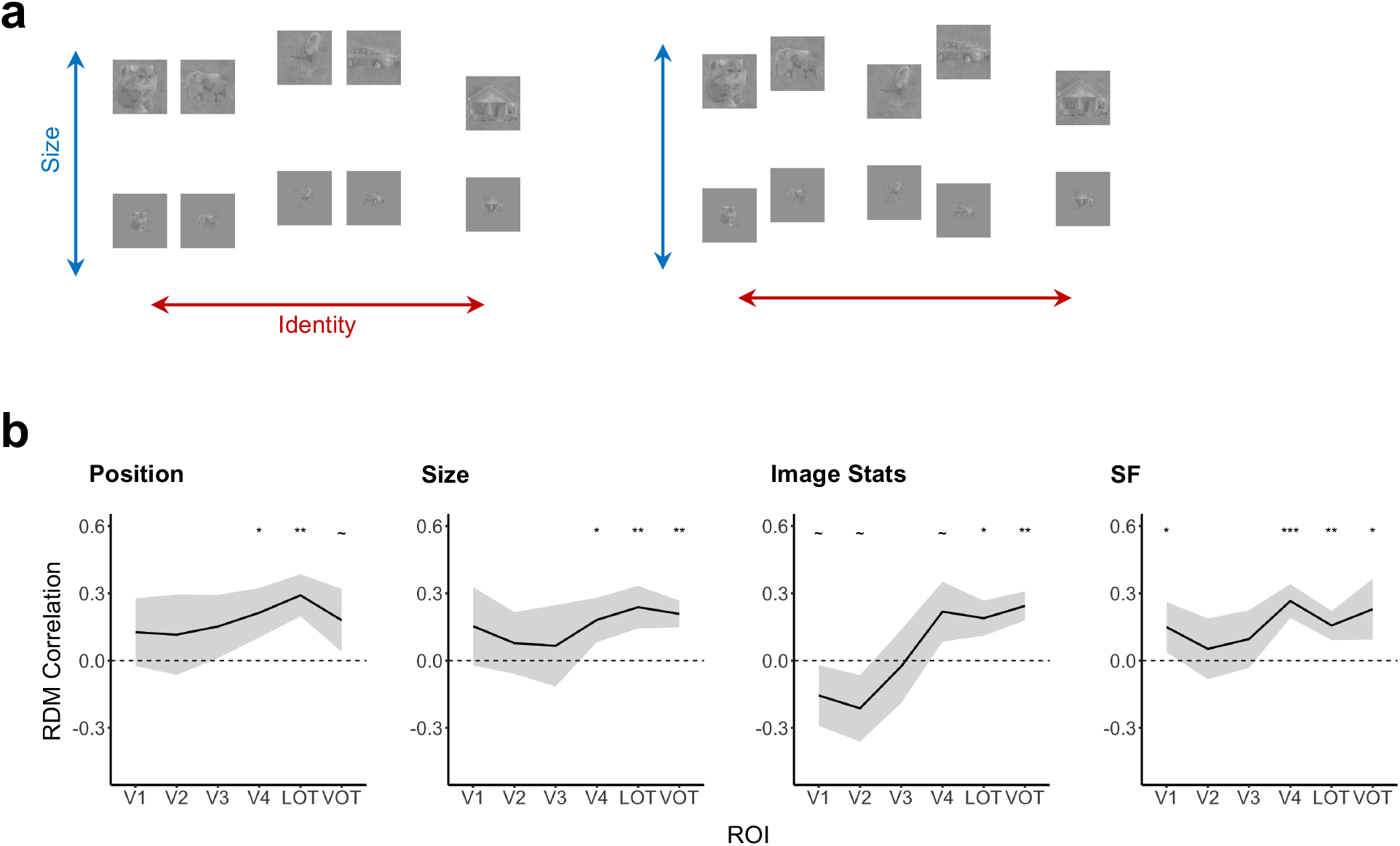
The effect of category similarity on pattern prediction generalizability. **a.** A schematic illustration of how object identity and non-identity features may be represented together in the high-dimensional neural representational space, using size as a non-identity feature. *Left panel*, coupled identity and non-identity representations. Categories that are closer in the identity dimension (e.g., the cat and elephant pair, and the chair and car pair) would also be similarly apart across the two states of the size transformation in the representational space. *Right panel*, decoupled identity and non-identity representations. Categories that are similar in the identity dimension are not necessarily similar across the two states of the size transformation. **b.** Correlations between the prediction similarity matrix and category similarity matrix for the 75 most reliable voxels included in each ROI. Results are plotted by transformation type showing the effect of category similarity on pattern prediction similarity as a function of the ROI. Significant values for pairwise t tests against 0 for each ROI are marked with asterisks at the top of each plot. All t tests were corrected for multiple comparison using the Benjamini-Hochberg method. ~ .05 < *p* < .10, * *p* <.05, ** *p* <.01, *** *p* <.001. Error bars represent the between-subjects 95% confidence interval.

To test if category similarity may increase pattern prediction, we first examined whether there was any positive correlation between the two. Overall, only the image stats transformation showed a significant main effect of ROI (*F*(5,29) = 9.28, *p* < 0.001; *Fs* < 2.00, *ps* > 0.1 for the other transformations), with V4, LOT and VOT showing greater correlations than V1, V2 and V3 in this transformation (*ts* > 2.50, *ps* < 0.05). Pairwise t-tests revealed greater than 0 correlation in LOT for all the transformations and in V4 and VOT for three out of the four transformations. However, only one lower visual region in one transformation showed a greater than 0 correlation (see the significance levels reported on Figure 6B for all the comparisons; additionally, see Figure 6-1 for similar results when all voxels from each ROI were included in analysis). Note that even when significant, these correlations were fairly modest, ranging from .15 to .25. Across the different ROIs, a positive linear trend from lower to higher visual regions was found for the image stats and SF transformations (*ts* > 2.81, *ps* < 0.05) but not for the other two transformations (*ts* < 1.19, *ps* > 0.13). Overall, these results showed that category similarity did increase pattern prediction to some extent, especially in higher visual regions; but the effect was overall fairly small.

## Discussion

Presently we still do not fully understand how object identity and non-identity information are simultaneously represented in the primate visual system. Building upon previous research efforts by Ward et al. (2018) and adapting their methodology, by analyzing data from two existing studies (Vaziri-Pashkam & Xu, 2019; Vaziri-Pashkam et al., 2019), here we conducted a comprehensive investigation to examine the existence of general mapping functions between fMRI responses to object categories in different states of non-identity transformations in the human brain. We examined and compared responses from human early visual areas V1 to V4 and two higher object processing regions, LOT and VOT to both Euclidean and non-Euclidean transformations of objects. For each transformation type and ROI, we derived a linear transformation matrix to link the fMRI response patterns of two states of an object category. Using this transformation matrix, we generated the predicted pattern of an object category in one state using its true pattern from the other state.

We first evaluated how well the predicted patterns of a category correlated with the true pattern of the same category. For all transformations, a linear mapping could capture a significant amount, but not all, of the variance of an object category’s fMRI pattern between two states of a transformation. Pattern predictions tended to be better for the categories included in the training data but did not benefit from including more categories in the training data. The linear mapping functions derived during training are thus not entirely category independent, but interact with the specific categories included in training. Higher visual regions did not exhibit better performance than lower visual regions.

In our second analysis, we evaluated whether the predicted patterns were closer to the true pattern of the same than different categories (i.e., pattern selectivity). Overall, the linear mapping predicted response patterns showed significant category selectivity, particularly for categories included in the training data across all ROIs and all transformations. For categories not included in the training data, only higher visual regions exhibited strong and consistent category selectivity when large numbers of categories were included in training. The linear mapping functions appeared to be tailored to a significant extent to the categories included in training.

As a third evaluation, we examined if the representational structure between categories was preserved for the predicted patterns. Linear mapping predicted response patterns were able to preserve the category representational structures in the true responses, particularly for categories included in the training data across all ROIs and all transformations. For categories not included in the training data, only higher visual regions were able to do so consistently; lower regions could do so only for the image stats transformation, but not for the other transformations. Predictions did not appear to be affected by the number of categories included in training. Across the different transformation types, a positive trend of better performance from lower to higher visual regions was only found in a little over half of the conditions tested, indicating that prediction was not always better in higher than lower regions.

Previous monkey neurophysiology and human fMRI studies have shown successful object decoding across different states of a transformation in higher level visual areas as well as successful non-identity feature decoding across different objects in both lower and higher level visual areas (e.g., Carlson et al., 2011; Cichy et al., 2011 & 2013; Grill-Spector et al., 1999; Hung et al., 2005; Hong et al., 2016; Rust & DiCarlo, 2010; Sawamura et al., 2005; Schwarzlose et al., 2008; Vaziri-Pashkam & Xu, 2019; Vaziri-Pashkam et al., 2019; Vuilleumier et al., 2002). These results suggest that identity and non-identity features are represented in the human visual system at least somewhat orthogonally. We found that the linear mapping functions derived were not entirely category independent, which could be due to either the near-orthogonal structure of the object representational space as depicted in Figure 1A, or data overfitting during training. In a near-orthogonal structure, identity and non-identity features can each be linearly decoded, but a learned linear mapping between two states of a transformation for one object category may not fully predict the neural response of a new object from one state to another.

We do not believe data overfitting played a significant role here. Had the object representational space contained category-independent orthogonal representations of the non-identity features, when more training data were included with increasing training set size, we would expect to see better prediction performance regardless of whether or not a category was included in the training data. However, we did not find any benefits of larger training set size for pattern prediction nor preservation of representational structure. For category selectivity, for categories included in the training data, performance tended to decrease with increasing training set sizes, but for categories not included in the training data, performance tended to increase with increasing training set sizes. Taken together, these results argue against a data overfitting account. Rather, they are more consistent with the object representational space being near-orthogonal, such that linear mapping of object responses between two states of a transformation may only partially capture how different object features are co-represented. To fully capture object responses between two states of a non-identity transformation, both a category independent component (as reported here) and a category dependent component would be needed.

It may be argued that, because the true pattern of an object category would always fluctuate somewhat due to noise, even if the true feature representational space is completely orthogonal, the measured feature space would always appear near-orthogonal. To account for such noise-driven pattern fluctuations, we used a split-half approach in the present study. Specifically, in all of our measures, we correlated the predicted patterns from one half of the data with the true pattern from the other half of the data and then assessed whether the correlation was comparable to the correlation of true patterns between the two halves of the data. Any fluctuation in response patterns would thus be captured and accounted for by the less than perfect correlation between the true patterns across the two halves of the data. Additionally, if the feature representational space was indeed completely orthogonal, because we tested on the left-out data not included in training, predictions should have been equally good for categories included in training and those that were not. However, we consistently found that predictions were in general better for categories included than those not included in training. Our approach was thus capable of detecting a completely orthogonal feature representational space if it existed in the data.

A feature-untangling view of object invariance may have predicted the existence of a stronger linear mapping in higher than lower visual regions as different object features become more separated during the course of visual information process in the ventral visual processing pathway (e.g., DiCarlo & Cox, 2007; Isik et al., 2013; Rolls, 2000; Rust & DiCarlo, 2010).

However, we did not find significant differences in prediction performance nor preservation of representational structure across ROIs. Although category selectivity was greater for higher than lower visual regions, this was likely due to object category representations being much stronger in higher than lower visual regions as revealed by previous fMRI decoding studies (e.g., Vaziri-Pashkam & Xu, 2017 & 2019; Vaziri-Pashkam et al., 2019). As such, greater category selectivity was expected in higher than lower visual regions even if linear mapping functions predicted responses equally well in both regions (see Figure 3 for a schematic explanation). Overall, the evidence shows that the near-orthogonal representation of object identity and non-identity features is present throughout the ventral visual regions. Thus, the non-identity features examined here are largely untangled from identity features much earlier on during visual processing in the human brain. It is therefore not the case that identity representation must be untangled from all non-identity features during ventral visual processing; but rather, such an untangling process may only apply to some of the non-identity features, such as viewpoint.

To further understand whether the near-orthogonal structure of the representational space co-varies with the similarity among the different object categories (i.e., if the ability of using the response of one category to predict that of another category would be determined by the similarity of these two categories in a given brain region), we tested the correlation between pattern prediction similarity and category similarity. Overall category similarity did increase pattern prediction to some extent, especially in higher visual regions; but the effect was fairly small.

In summary, using a representational transformation analysis, we show that the entire human ventral visual system can link object responses in different states of non-identity transformations through linear mapping functions for both Euclidean and non-Euclidean transformations. These mapping functions are not entirely identity-independent, suggesting that object identity and non-identity features are represented in a near, rather than completely, orthogonal manner. Our study provides a useful framework to more precisely characterize how different types of object features may be represented together during visual processing in the human brain.

## Acknowledgements

We thank members of Visual Cognitive Neuroscience Lab, Turk-Browne Lab, Holmes Lab, and Yale Cognitive and Neural Computation Lab for their helpful feedback on this project. This research was supported by the National Institute of Health Grants (1R01EY030854 and 1R01EY022355) to Y.X.

---

**Figure 2-1.**
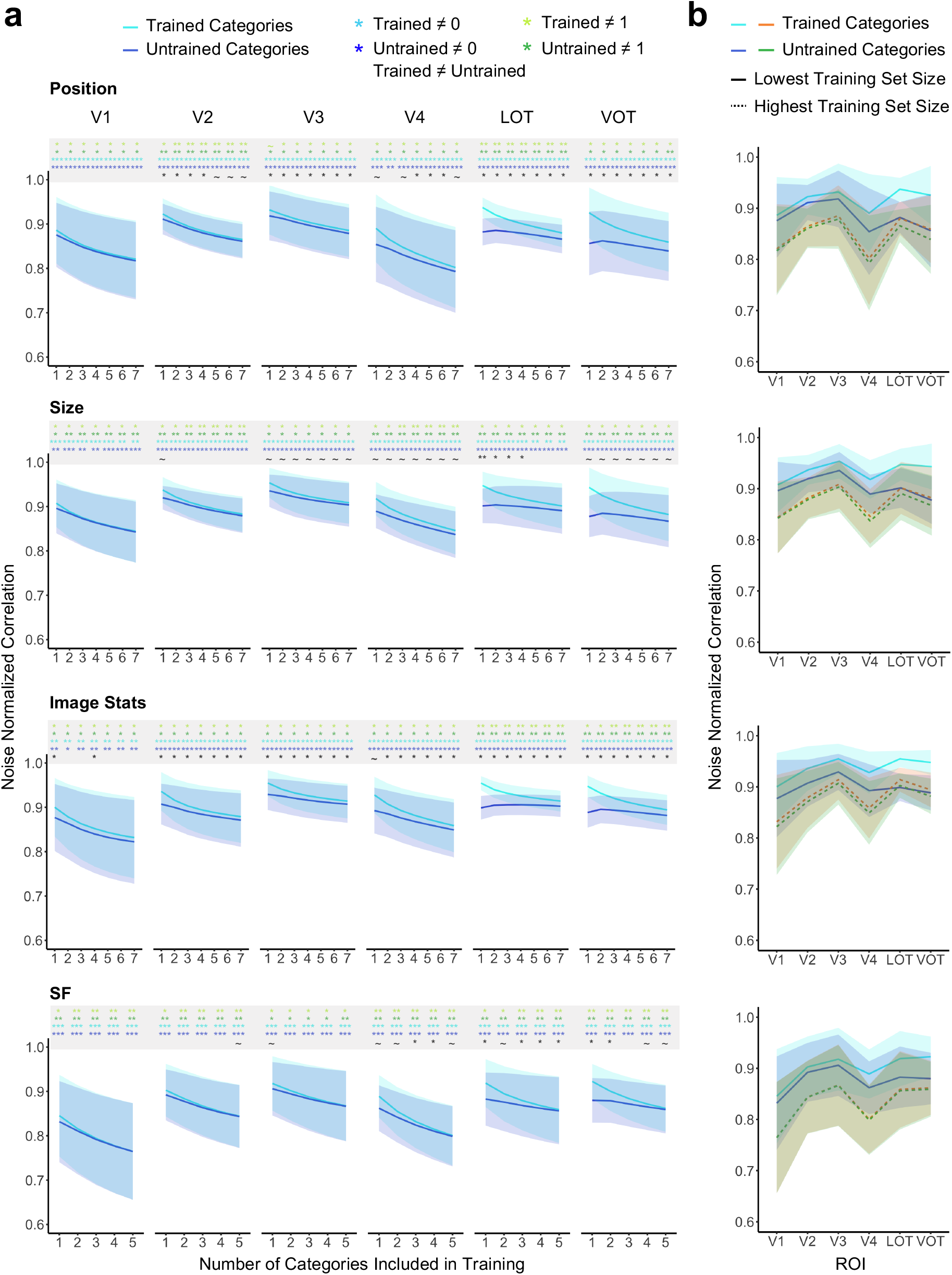
Correlations between the predicted and the corresponding true patterns for each transformation for all the voxels included in each ROI. The correlations were normalized by the reliability of each ROI (see Methods). **a.** Results plotted by ROI and transformation type showing pattern prediction for categories included in the training data (Trained Categories) and those that were not (Untrained Categories) as a function of the number of categories included in the training data. Significant values for pairwise t tests against 0 and 1 for each condition and for the difference between the trained and untrained categories for each training set size are marked with asterisks at the top of each plot. All t tests were corrected for multiple comparison using the Benjamini-Hochberg method. ~ .05 < *p* < .10, * *p* <.05, ** *p* <.01, *** *p* <.001. **b.** Results plotted by transformation type showing pattern prediction as a function of the ROI including only the smallest and the largest training set sizes. Error bars represent the between-subjects 95% confidence interval.

**Figure 4-1.**
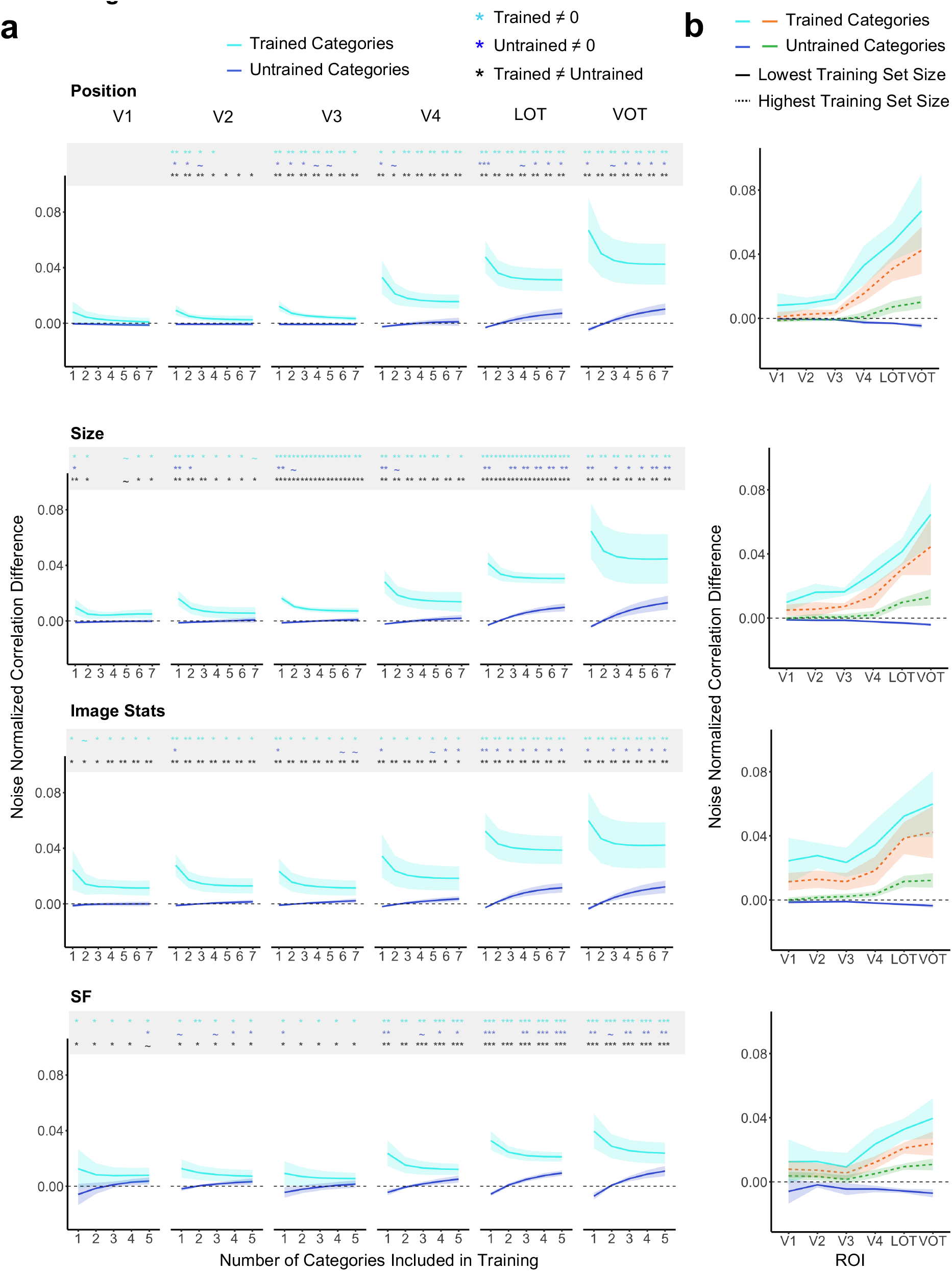
Category selectivity for the predicted patterns for all the voxels included in each ROI. Category selectivity was calculated as the correlation difference between the predicted and true patterns of the same category and that of different categories. **a.** Results plotted by ROI and transformation type showing category selectivity for categories included in the training data (Trained Categories) and those that were not (Untrained Categories) as a function of the number of categories included in the training data. Significant values for pairwise t tests against 0 for each condition and for the difference between the trained and untrained categories for each training set size are marked with asterisks at the top of each plot. All t tests were corrected for multiple comparison using the Benjamini-Hochberg method. ~ .05 < *p* < .10, * *p* <.05, ** *p* <.01, *** *p* <.001. **b.** Results plotted by transformation type showing category selectivity as a function of the ROI including only the smallest and the largest training set sizes. Error bars represent the between-subjects 95% confidence interval.

**Figure 5-1.**
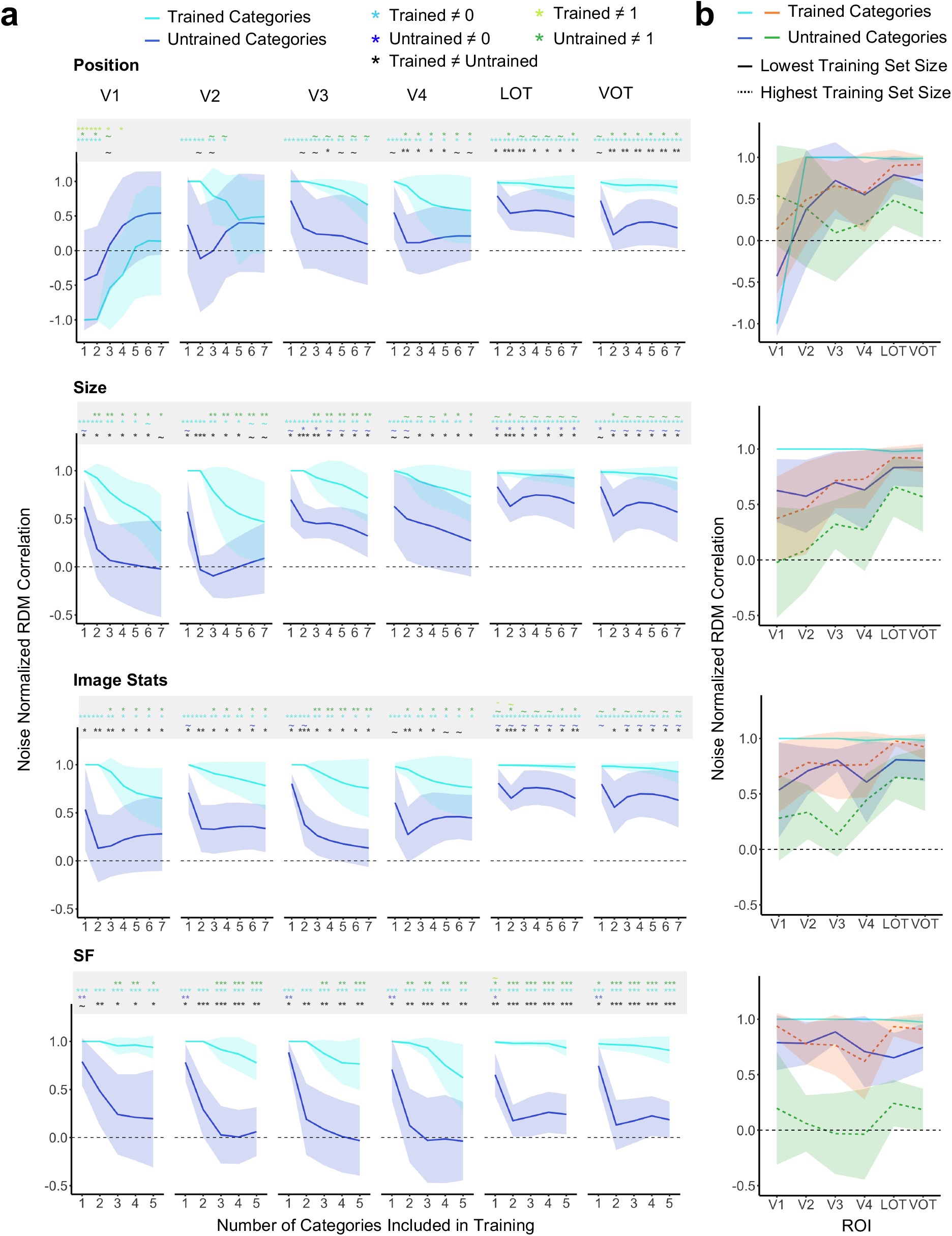
Category RDM correlation between the predicted and true patterns for all the voxels included in each ROI. The correlations were normalized by the reliability of each ROI (see Methods). **a.** Results plotted by ROI and transformation type showing category structure preservation for categories included in the training data (Trained Categories) and those that were not (Untrained Categories) as a function of the number of categories included in the training data. Significant values for pairwise t tests against 0 and 1 for each condition and for the difference between the trained and untrained categories for each training set size are marked with asterisks at the top of each plot. All t tests were corrected for multiple comparison using the Benjamini-Hochberg method. ~ .05 < *p* < .10, * *p* <.05, ** *p* <.01, *** *p* <.001. **b.** Results plotted by transformation type showing category structure preservation as a function of the ROI including only the smallest and the largest training set sizes. Error bars represent the between-subjects 95% confidence interval.

**Figure 6-1.**
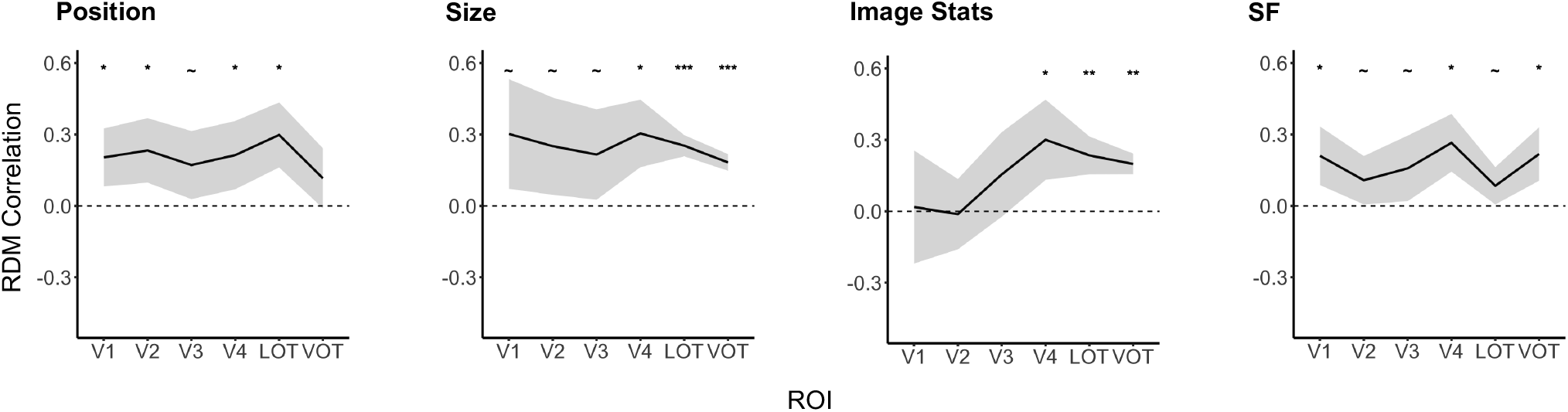
Correlations between the prediction similarity matrix and category similarity matrix for all the voxels included in each ROI. Results are plotted by transformation type showing the effect of category similarity on pattern prediction similarity as a function of the ROI. Significant values for pairwise t tests against 0 for each ROI are marked with asterisks at the top of each plot. All t tests were corrected for multiple comparison using the Benjamini-Hochberg method. ~ .05 < *p* < .10, * *p* <.05, ** *p* <.01, *** *p* <.001. Error bars represent the between-subjects 95% confidence interval.

## References

Benjamini, Y., & Hochberg, Y. (1995). Controlling the False Discovery Rate: A Practical and Powerful Approach to Multiple Testing. Journal of the Royal Statistical Society. Series B (Methodological), 57(1), 289–300. JSTOR.

Carlson, T. A., Hogendoorn, H., Kanai, R., Mesik, J., & Turret, J. (2011). High temporal resolution decoding of object position and category. Journal of Vision, 11(10), 9–9. https://doi.org/10.1167/11.10.9

Cichy, R. M., Chen, Y., & Haynes, J.-D. (2011). Encoding the identity and location of objects in human LOC. NeuroImage, 54(3), 2297–2307. https://doi.org/10.1016/j.neuroimage.2010.09.044

Cichy, R. M., Sterzer, P., Heinzle, J., Elliott, L. T., Ramirez, F., & Haynes, J.-D. (2013). Probing principles of large-scale object representation: Category preference and location encoding. Human Brain Mapping, 34(7), 1636–1651. https://doi.org/10.1002/hbm.22020

DiCarlo, J. J., & Cox, D. D. (2007). Untangling invariant object recognition. Trends in Cognitive Sciences, 11(8), 333–341. https://doi.org/10.1016/j.tics.2007.06.010

Grill-Spector, K., Kushnir, T., Hendler, T., Edelman, S., Itzchak, Y., & Malach, R. (1998). A sequence of object-processing stages revealed by fMRI in the human occipital lobe. Human Brain Mapping, 6(4), 316–328. https://doi.org/10.1002/(SICI)1097-0193(1998)6:4<316::AID-HBM9>3.0.CO;2-6

Grill-Spector, K., Kushnir, T., Edelman, S., Avidan, G., Itzchak, Y., & Malach, R. (1999). Differential Processing of Objects under Various Viewing Conditions in the Human Lateral Occipital Complex. Neuron, 24(1), 187–203. https://doi.org/10.1016/S0896-6273(00)80832-6

Haxby, J. V., Gobbini, M. I., Furey, M. L., Ishai, A., Schouten, J. L., & Pietrini, P. (2001). Distributed and Overlapping Representations of Faces and Objects in Ventral Temporal Cortex. Science, 293(5539), 2425–2430. https://doi.org/10.1126/science.1063736

Hong, H., Yamins, D. L. K., Majaj, N. J., & DiCarlo, J. J. (2016). Explicit information for category-orthogonal object properties increases along the ventral stream. Nature Neuroscience, 19(4), 613–622. https://doi.org/10.1038/nn.4247

Hung, C. P., Kreiman, G., Poggio, T., & DiCarlo, J. J. (2005). Fast Readout of Object Identity from Macaque Inferior Temporal Cortex. Science, 310(5749), 863–866. https://doi.org/10.1126/science.1117593

Isik, L., Meyers, E. M., Leibo, J. Z., & Poggio, T. (2013). The dynamics of invariant object recognition in the human visual system. Journal of Neurophysiology, 111(1), 91–102. https://doi.org/10.1152/jn.00394.2013

Ito, M., Tamura, H., Fujita, I., & Tanaka, K. (1995). Size and position invariance of neuronal responses in monkey inferotemporal cortex. Journal of Neurophysiology, 73(1), 218–226. https://doi.org/10.1152/jn.1995.73.1.218

Kamitani, Y., & Tong, F. (2005). Decoding the visual and subjective contents of the human brain. Nature Neuroscience, 8(5), 679–685. https://doi.org/10.1038/nn1444

Kietzmann, T. C., Spoerer, C. J., Sörensen, L. K. A., Cichy, R. M., Hauk, O., & Kriegeskorte, N. (2019). Recurrence is required to capture the representational dynamics of the human visual system. Proceedings of the National Academy of Sciences, 116(43), 21854–21863. https://doi.org/10.1073/pnas.1905544116

Kourtzi, Z., & Kanwisher, N. (2000). Cortical Regions Involved in Perceiving Object Shape. Journal of Neuroscience, 20(9), 3310–3318. https://doi.org/10.1523/JNEUROSCI.20-09-03310.2000

Kriegeskorte, N., & Kievit, R. A. (2013). Representational geometry: Integrating cognition, computation, and the brain. Trends in Cognitive Sciences, 17(8), 401–412. https://doi.org/10.1016/j.tics.2013.06.007

Kriegeskorte, N., Mur, M., Ruff, D. A., Kiani, R., Bodurka, J., Esteky, H., Tanaka, K., & Bandettini, P. A. (2008). Matching Categorical Object Representations in Inferior Temporal Cortex of Man and Monkey. Neuron, 60(6), 1126–1141. https://doi.org/10.1016/j.neuron.2008.10.043

Logothetis, N. K., Pauls, J., & Poggio, T. (1995). Shape representation in the inferior temporal cortex of monkeys. Current Biology, 5(5), 552–563. https://doi.org/10.1016/S0960-9822(95)00108-4

Malach, R., Reppas, J. B., Benson, R. R., Kwong, K. K., Jiang, H., Kennedy, W. A., Ledden, P. J., Brady, T. J., Rosen, B. R., & Tootell, R. B. (1995). Object-related activity revealed by functional magnetic resonance imaging in human occipital cortex. Proceedings of the National Academy of Sciences, 92(18), 8135–8139. https://doi.org/10.1073/pnas.92.18.8135

R Core Team (2018) R: a language and environment for statistical computing. Vienna: R Foundation for Statistical Computing. Available at http://www.R-project.org/.

Rolls, E. T. (2000). Functions of the Primate Temporal Lobe Cortical Visual Areas in Invariant Visual Object and Face Recognition. Neuron, 27(2), 205–218. https://doi.org/10.1016/S0896-6273(00)00030-1

Rust, N. C., & DiCarlo, J. J. (2010). Selectivity and Tolerance (“Invariance”) Both Increase as Visual Information Propagates from Cortical Area V4 to IT. Journal of Neuroscience, 30(39), 12978–12995. https://doi.org/10.1523/JNEUROSCI.0179-10.2010

Sawamura, H., Georgieva, S., Vogels, R., Vanduffel, W., & Orban, G. A. (2005). Using Functional Magnetic Resonance Imaging to Assess Adaptation and Size Invariance of Shape Processing by Humans and Monkeys. Journal of Neuroscience, 25(17), 4294–4306. https://doi.org/10.1523/JNEUROSCI.0377-05.2005

Schwarzlose, R. F., Swisher, J. D., Dang, S., & Kanwisher, N. (2008). The distribution of category and location information across object-selective regions in human visual cortex. Proceedings of the National Academy of Sciences, 105(11), 4447–4452. https://doi.org/10.1073/pnas.0800431105

Tarhan, L., & Konkle, T. (2019). Reliability-based voxel selection. NeuroImage, 116350. https://doi.org/10.1016/j.neuroimage.2019.116350

Vaziri-Pashkam, M., & Xu, Y. (2017). Goal-Directed Visual Processing Differentially Impacts Human Ventral and Dorsal Visual Representations. The Journal of Neuroscience, 37(36), 8767–8782. https://doi.org/10.1523/JNEUROSCI.3392-16.2017

Vaziri-Pashkam, M., Taylor, J., & Xu, Y. (2019). Spatial Frequency Tolerant Visual Object Representations in the Human Ventral and Dorsal Visual Processing Pathways. Journal of Cognitive Neuroscience, 31(1), 49–63. https://doi.org/10.1162/jocn_a_01335

Vaziri-Pashkam, M., & Xu, Y. (2019). An Information-Driven 2-Pathway Characterization of Occipitotemporal and Posterior Parietal Visual Object Representations. Cerebral Cortex, 29(5), 2034–2050. https://doi.org/10.1093/cercor/bhy080

Vuilleumier, P., Henson, R. N., Driver, J., & Dolan, R. J. (2002). Multiple levels of visual object constancy revealed by event-related fMRI of repetition priming. Nature Neuroscience, 5(5), 491–499. https://doi.org/10.1038/nn839

Ward, E. J., Isik, L., & Chun, M. M. (2018). General Transformations of Object Representations in Human Visual Cortex. The Journal of Neuroscience, 38(40), 8526–8537. https://doi.org/10.1523/JNEUROSCI.2800-17.2018

Willenbockel, V., Sadr, J., Fiset, D., Horne, G. O., Gosselin, F., & Tanaka, J. W. (2010). Controlling low-level image properties: The SHINE toolbox. Behavior Research Methods, 42(3), 671–684. https://doi.org/10.3758/BRM.42.3.671

